# Potential of host serum protein biosignatures in the diagnosis of tuberculous meningitis in children

**DOI:** 10.1101/670323

**Authors:** Charles M Manyelo, Regan S Solomons, Candice I Snyders, Kim Stanley, Gerhard Walzl, Novel N Chegou

**Affiliations:** DST-NRF Centre of Excellence for Biomedical Tuberculosis Research; South African Medical Research Council Centre for Tuberculosis Research; Division of Molecular Biology and Human Genetics, Faculty of Medicine and Health Sciences, Stellenbosch University; Department of Paediatrics and Child Health, Faculty of Medicine and Health Sciences, Stellenbosch University, Cape Town, South Africa

**Author notes:** Contributed equally (Charles M Manyelo, Regan S Solomons).

**Keywords:** Tuberculosis, meningitis, biomarkers, sensitivity, specificity, diagnosis, child

## Abstract

**Background:** Tuberculous meningitis (TBM) is the most severe form of tuberculosis and results in high morbidity and mortality in children. Diagnostic delay contributes to the poor outcome. There is an urgent need for new tools for the rapid diagnosis of TBM, especially in children.

**Methods:** We collected serum samples from children in whom TBM was suspected at a tertiary hospital in Cape Town, South Africa. Children were subsequently classified as having TBM or no TBM using a published uniform research case-definition. Using a multiplex cytokine array platform, we investigated the concentrations of serum biomarkers comprising 7-markers that were previously found to be of value in the diagnosis of adult pulmonary TB (CRP, SAA, CFH, IFN-γ, IP-10, Apo-AI and transthyretin) plus other potentially useful host biomarkers as diagnostic candidates for TBM.

**Findings:** Of 47 children included in the study, 23 (48.9%) had a final diagnosis of TBM of which six had HIV co-infection. A modified version of the adult 7-marker biosignature in which transthyretin was replaced by NCAM1, diagnosed TBM in children with AUC of 0.80 (95% CI, 0.67-0.92), sensitivity of 73.9% (95% CI, 51.6-89.8%) and specificity of 66.7% (95% CI, 44.7-84.4%). A new childhood TBM specific 3-marker biosignature (adipsin, Aβ42 and IL-10) showed potential in the diagnosis of TBM, with AUC of 0.84 (95% CI, 0.73-0.96), sensitivity of 82.6% (95 CI, 61.2-95.0%) and specificity of 75.0% (95% CI, 53.3-90.2%) after leave-one-out cross validation.

**Conclusion:** An adult 7-marker serum protein biosignature showed potential in the diagnosis of TBM in children. However, a smaller childhood TBM-specific biosignature demonstrated improved performance characteristics. Our data indicates that blood-based biomarkers may be useful in the diagnosis of childhood TBM and require further investigation.

## Introduction

Globally, tuberculosis (TB) is currently one of the top 9 causes of death, ranking above HIV and malaria (1). According to the World Health Organisation (WHO) 2018 TB report, 10% of all reported TB cases in 2017 were in children (2), and 210 000 children died from TB (1). About 20-25% of paediatric TB cases are extra pulmonary TB (EPTB), including tuberculous meningitis (TBM) (3). In a clinical and laboratory surveillance study over a one year period in the Western Cape Province of South Africa almost one-fifth of children with TB had a disseminated form (TB meningitis or miliary TB) (4).

TBM is a central nervous system infection caused by Mycobacterium tuberculosis (M.tb) and it is the most severe form of tuberculosis, with high morbidity and mortality (4, 5). The rate of TBM is higher among children than adults (7). A study in South Africa reported that TBM is the most common form of bacterial meningitis among children (8). Outcomes of TBM in children include death in up to 50% of cases and neurological sequelae in up to 53.9% of survivors (9, 10). The poor outcomes of TBM are mainly due to delayed diagnosis and late initiation of anti-tuberculosis therapy (11). The improvement of case detection and early administration of curative treatment are key in reducing high morbidity and mortality associated with paediatric tuberculosis (12). However, the diagnosis of TBM in children is challenging, due to sub-optimal performance of the currently available laboratory diagnostic methods. Smear microscopy remains the most widely used test for bacteriological confirmation of TB in clinical specimens, particularly in resource-constrained settings (13). The main limitation of this test is poor sensitivity due to its requirement of 5000 to 10000 M.tb bacilli per ml of sample for a positive result. In diagnosing tuberculous meningitis, smear microscopy has a poor sensitivity of about 10-20% (9). M.tb culture of biological fluids remains the gold standard test for diagnosing TB disease. Although culture has a relatively highly sensitivity (60-70%) for diagnosing TBM (in comparison to smear microscopy), the turnaround time for culture is up to 42 days (9). Furthermore, culture is expensive, prone to contamination and requires extensive laboratory infrastructure, which is not often available in resource-constrained settings (14). The GeneXpert MTB/RIF test®, arguably the most important recent advance in TB diagnosis, yields results within 2 hours and detects resistance to rifampicin as a proxy for the presence of MDR strains. However, GeneXpert has several well-publicised shortcomings including low negative predictive value, cost effectiveness and the requirement for technical infrastructure (10, 12).

GeneXpert cannot exclude tuberculous meningitis due to its imperfect sensitivity and negative predictive value (16). A study conducted in Uganda reported sensitivity of 28% when using 2 mL of uncentrifuged CSF, and the sensitivity improved to 72% when a large volume (6mL) of centrifuged CSF was used (17), suggesting that high volumes of CSF are required to obtain a positive TBM diagnosis using Xpert. GeneXpert Ultra (the next generation assay) has been shown to overcome some of the shortcomings of GeneXpert MTB/RIF as it showed improved sensitivity of 95% and negative predictive value of 99% in a more recent study on HIV positive adults with TBM (16). However, concerns over lowered specificity and positive predictive value compared to GeneXpert MTB/RIF need to be allayed by prospective testing in a large population including children and those that are HIV-uninfected (16, 18, 19). The main common limitation to the above-described tests is the difficulty in obtaining the diagnostic sample (CSF) due to the need for more invasive sampling methods and low yields of mycobacteria obtained from childhood CSF samples, as a result of the paucibacillary nature of the disease.

The diagnosis of TBM is mostly based on a combination of suggestive clinical findings, multiple suggestive laboratory tests on the CSF, supportive features on brain imaging findings and the exclusion of other possible causes of meningitis. These criteria are unreliable as individual tests, and often require referral of children to tertiary healthcare facilities for the performance of advanced testing, due to the unavailability of most of the relevant techniques in primary and secondary healthcare facilities, leading to further delays in the initiation of treatment (20). Consequently, children seen at primary and secondary healthcare facilities often have multiple missed opportunities; up to six visits before eventual diagnosis of TBM is made (20). There is an urgent need for new TBM diagnostic tools suitable for use in children at the point of care or bedside.

Host biomarker-based tests may be valuable in the diagnosis of TBM as they have previously been shown to be potentially useful in other extrapulmonary forms of TB (21), and may be easily converted into point-of-care or bedside tests (20, 21). Most studies investigating host biomarkers for diagnosis of neurological disorders have focused on CSF as a biological specimen from the site of infection (24), including in the diagnosis of TBM (23, 24). Despite the potential of CSF-based host biomarkers in the diagnosis of TBM, the collection of CSF requires lumbar puncture, an invasive procedure which requires special skill and training and might not be readily available, leading to delay in diagnosis.

As several studies have shown the potential of blood-based host protein biomarkers in the diagnosis of TB disease, albeit mostly adult pulmonary TB-based studies thus far (25, 26), we hypothesised that blood-based host protein biosignatures may be useful in the diagnosis of TBM in children. Obtaining serum is less invasive when compared to obtaining CSF. A test based on serum host proteins may be more easily applicable in resource-limited settings due to ease of sample collection. Furthermore, serum based tests may be easily converted to finger prick-based assays as is currently being done elsewhere (www.screen-tb.eu). The aim of the present study was to ascertain whether host biomarkers that have shown potential in the diagnosis of adult pulmonary TB in serum and plasma samples (25, 26) possessed diagnostic potential for childhood TBM. We were specifically interested in evaluating the performance of a previously established adult seven-marker serum protein biosignature (CRP, transthyretin, IFN-γ, CFH, Apo-AI, IP-10, and SAA) (28) as a tool for the diagnosis of childhood TBM, and to also evaluate the potential of other host biomarkers.

## Materials and Methods

### Study participants

Participants enrolled into the present study were children who presented with signs and symptoms suggestive of meningitis and requiring CSF examination for routine diagnostic purposes at the Tygerberg Academic Hospital in Cape Town, South Africa between November 2016 and November 2017 as previously described (29). Children were eligible for participation in the study if they were between the ages of 3 months and 13 years, provided that written informed consent was obtained from the parents or legal guardians. Assent was obtained from children older than 7 years if they had a normal level of consciousness i.e., a Glasgow Coma Score (GCS) of 15/15. The study was approved by the Health Research Ethics Committee of the University of Stellenbosch, Tygerberg Academic Hospital, and the Western Cape Provincial Government.

After collection of specimens for routine diagnostic purposes, an additional 1ml of blood was collected into a BD Vacutainer® serum tube and processed within an average of 2 hours from collection, for the purposes of the present study. Blood samples were centrifuged at 1200 xg for 10 minutes, followed by aliquoting of serum and storage at −80°C until analyzed.

### Diagnostic assessment

As previously described (29), a comprehensive clinical examination was performed on all patients by a specialist paediatric neurologist. Following routine clinical investigations, computed tomography (CT) of the brain, air-encephalography, and magnetic resonance (MR) imaging were performed as clinically indicated. CSF was obtained through lumbar puncture and investigations including appearance and colour determination, differential cell counts, CSF protein and glucose levels and other routinely investigated parameters were done, followed by centrifugation, Gram staining, India ink examination, culture of the centrifuged sediment on blood agar plates (for pyogenic bacteria), Auramine “O” staining and fluorescence microscopy, culture using the mycobacterium growth indicator tubes (MGIT) ™ (Becton and Dickinson) and examination for M.tb DNA using the HAIN Genotype MTBDRplus kit (Hain Lifescience GmbH, Germany). All data generated from the study were recorded in a study specific RedCap web-based database.

### Classification of study participants

Patients were classified as probable or definite TBM according to a uniform research case definition (30) based on a scoring system consisting of clinical, CSF, neuroimaging findings and evidence of extraneural TB. TBM was classified as ‘probable’ when patients scored ≥12 when neuroimaging was available and ≥10 when neuroimaging was unavailable. In this cohort, a diagnosis of definite TBM was made if acid-fast bacilli were present in the CSF, M.tb was isolated from the CSF by culture, or nucleic acid amplification test of CSF yielded positive results. For the purposes of this study, the TBM group included probable TBM and definite TBM, whereas the no-TBM group included children with other types of meningitis including bacterial and viral meningitis, and a wide range of other diagnoses (namely: asphyxia, autoimmune encephalitis, febrile seizure, Guilain Barre, HIV encephalopathy, focal seizures, leukemia, miliary TB (with lymyphocytic interstitial pneumonitis), developmental delay, breakthrough seizure, gastroenteritis (caused by shock), idiopathic intracranial hypertention (IIH), viral gastroenteritis (Adeno and Rota) and encephalopathy, stroke, mitochondrial diagnosis, viral pneumonia, febrile seizure and acute gastroenteritis) as mentioned in Table 1 and also previously reported (29).

**Table 1:** Clinical and demographic characteristics of children included in the study

|  | **All** | **TBM** | **No-TBM<sup>#</sup>** |
| --- | --- | --- | --- |
| Number of participants | 47 | 23 | 24 |
| Mean age, months± SD | 41.6 ± 41.5 | 31.5±34.8 | 51.3±45.7 |
| Males, n (%) | 30(63.8) | 13(56.5) | 17(70.8) |
| HIV Positive, n/no tested | 6 /37 | 0 /22 | 6 /15 |

<sup>#</sup>The no-TBM group included children with bacterial meningitis (n=2), viral meningitis (n=2), asphyxia (n=1), autoimmune encephalitis (n=1), febrile seizure (n=3), Guillain Barre (n=1), HIV encephalopathy (n=1), focal seizures (n=1), leukemia (n=1), Miliary TB (With lymphocytic interstitial pneumonitis) (n=1), Developmental delay (n=1), Breakthrough seizure (n=1), Gastroenteritis (Caused by shock) (n=1), Idiopathic intracranial hypertension (IIH) (n=1), Viral Gastroenteritis (Adeno and Rota) and encephalopathy (n=1), Stroke (n=1), Mitochondrial diagnosis (n=1), viral pneumonia (This included also SAM and nosocomial sepsis) (n=1), Febrile Seizure and Acute gastroenteritis (n=1) and other (n=1). Adapted from (29).

### Immunoassays

We evaluated the concentrations of biomarkers assessed in our previous study (29) in serum samples obtained from the same study participants. As reported in our previous study, the concentrations of 69 host biomarkers including six of the proteins comprising the previously established adult seven-marker serum protein biosignature (CRP, SAA, complement factor H, IFN-γ, IP-10 and Apo AI) (26), were evaluated by ELISA (cathelicidin LL-37) or using the Luminex platform (all other biomarkers). Transthyretin, one of the biomarkers comprising the previously established adult seven-marker signature was not assessed due to discontinued supply from the manufacturer.

Serum cathelicidin LL-37 levels were evaluated using an ELISA kit purchased from Elabscience Biotechnology Inc. (Catalog #E-EL-H2438). Experiments were done according to the procedure recommended by the manufacturer after which optical densities (OD) were read at 450nm by an automated microplate reader (iMark™ Microplate Reader, Bio Rad Laboratories). The mean OD of the blank wells was subtracted from the OD of the sample wells and the background-corrected ODs used for statistical analysis.

Serum levels of CCL1(I-309), CCL2(MCP-1), CCL3(MIP-1α), CCL4(MIP-1β), CD40 ligand (CD40L), CXCL8(IL-8), CXCL9(MIG), CXCL10(IP-10), granulocyte colony-stimulating factor (G-CSF), granulocyte-macrophage colony-stimulating factor(GM-CSF), interferon (IFN)-γ, interleukin (IL)-1β, IL-10, IL-12/23(p40), IL-17A, IL-13, IL-21, IL-4, IL-6, IL-7, matrix metalloproteinase (MMP)-1, MMP-8, transforming growth factor (TGF)-α, tumour necrosis factor (TNF)-α, soluble neural cell adhesion molecule (sNCAM-1/CD56), MMP-7, VEGF-A, ferritin and MMP-9 were assessed in Luminex kits purchased from R&D Systems Inc. (Bio-Techne), Minneapolis, USA wherease those of apolipoprotein (Apo)-AI, Apo-CIII, complement C3, complement factor H, BDNF, cathepsin D, soluble intracellular adhesion molecule (sICAM)-1, myeloperoxidase (MPO), platelet derived growth factor (PDGF)-AA, CCL5(RANTES), PDGF-AB/BB, soluble vascular adhesion molecule (sVCAM-1), plasminogen activator inhibitor (PAI)-1(total), S100 calcium-binding protein B (S100B), amyloid beta 1-40 (Ab40), Ab42, soluble receptor for advanced glycation end products (sRAGE), Glial cell-derived neurotrophic factor (GDNF), C reactive protein (CRP), alpha-2-antitrypsin (A1AT), pigment epithelium-derived factor (PEDF), serum amyloid P (SAP), CCL18(MIP-4/PARC), complement C4 (CC4), CC2, CC4b, CC5, CC5a, CC9, complement factor D (adipsin/CFD), mannose binding lectin (MBL), complement factor 1 (CF1), sP-selectin, von Willebrand factor-cleaving protease (ADAMTS13), D-DIMER, growth differentiation factor (GDF)-15, myoglobin, lipocalin2 (NGAL), and serum amyloid A (SAA) were evaluated in kits purchased form Merck Millipore, Billerica, MA, USA.

All biomarkers were evaluated following the instructions of the respective kit manufacturers (R&D Systems and Merck Millipore, respectively) in a blinded manner. All experiments were performed on the Bio Plex platform (Bio Rad Laboratories, Hercules, USA) in an ISO15189 accredited laboratory. Data acquisition and analysis of median fluorescent intensity was done using the Bio Plex Manager Version 6.1 software (Bio Rad Laboratories). The values of analytes in the quality control reagents evaluated with the samples were within their expected ranges.

### Statistical analysis

Data were analysed using Statistica (TIBCO Software Inc., CA, USA), and GraphPad Prism version 6 (Graphpad software, CA, USA). The Mann Whitney U test was used to compare the differences in the concentrations of host biomarkers between the TBM and the no-TBM groups. The receiver operator characteristics (ROC) curve analysis procedure was used to assess the diagnostic accuracy of individual host biomarkers for TBM. Optimal cut-off values and associated sensitivities and specificities were selected based on the Youden’s index (31). The utility of combinations of biomarkers in the diagnosis of TBM was ascertained by general discriminant analysis (GDA), followed by leave-one-out cross validation. Data were log-transformed to prior to GDA.

## Results

A total of 47 children who presented with signs and symptoms strongly suggestive of TBM were included in the study, 30 (63.8%) of whom were males. The mean age of all the children was 41.6 ± 41.5 months and six out of 37 with known HIV status (16.2%) were HIV infected. Using a composite reference standard based on a uniform research case definition of TBM (29), 23 of the children were diagnosed with definite (n=2) or probable (n=21) TBM. The 24 children without TBM included children with bacterial meningitis (n=2), viral meningitis (n=2) and children with other diagnoses as mentioned in Table 1 and also previously reported (29). (Table 1, Figure 1).

**Figure 1:**
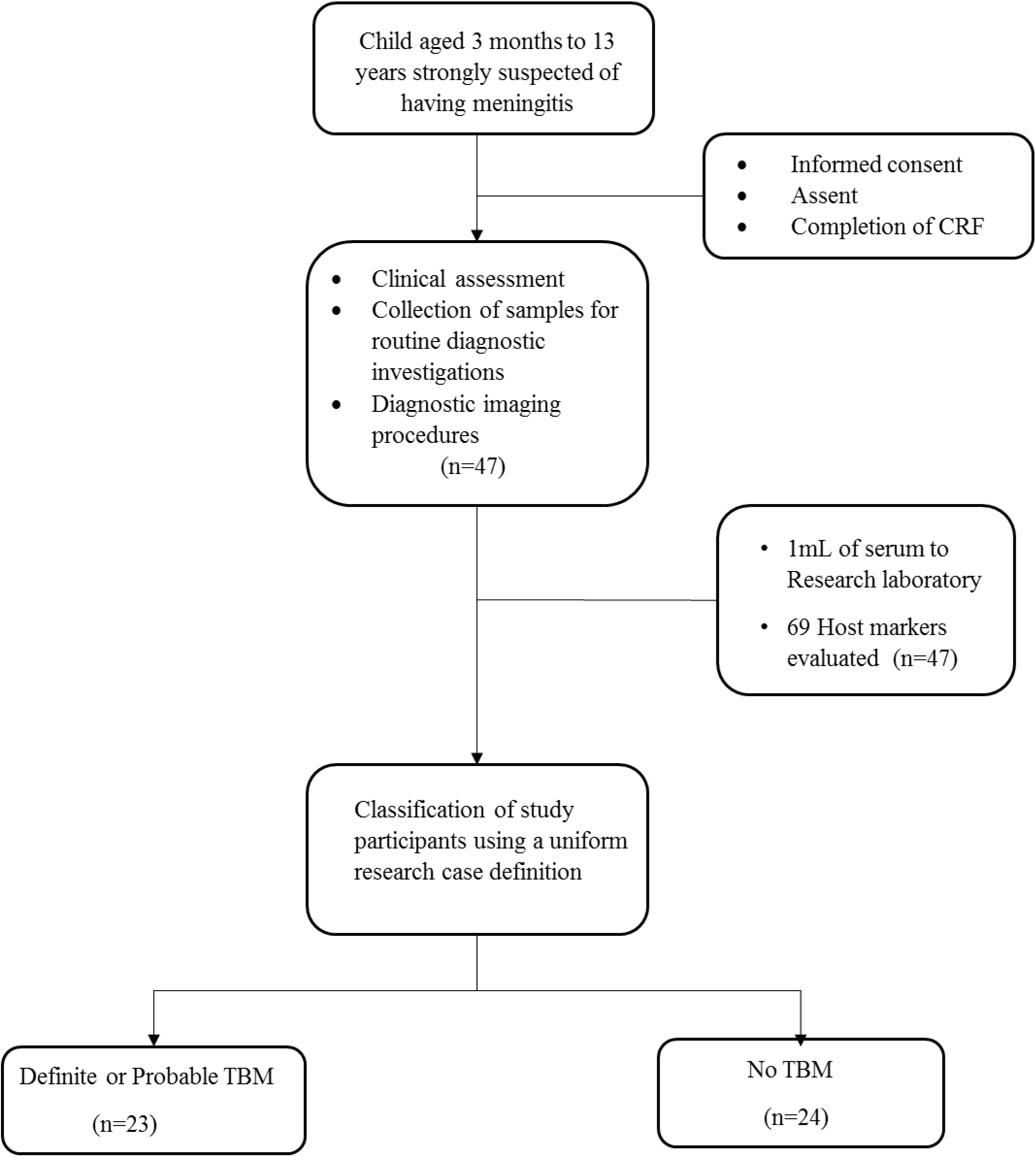
Flow diagram showing the study design and classification of study participants. CRF, case report form; TBM, Tuberculous meningitis; No-TBM, Individuals presenting with symptoms and investigated for TB but TBM ruled out. The No TBM group included bacterial meningitis (n=2), viral meningitis (n=2) and children with other diagnoses (Table 1). Adapted from Manyelo et al, 2019, Mediators of Inflammation, In Press.

### Application of the previously established adult 7-marker serum protein biosignature in the diagnosis of TBM in children

When the concentrations of the six available markers, from the adult pulmonary TB seven-marker signature (CRP, IFN-γ, IP-10, CFH, Apo-A1 and SAA) were evaluated in serum samples from children with TBM versus those without TBM individually, significant differences were obtained for CFH only. After ROC curve analysis, the most useful individual marker from this signature, as determined by AUC was CFH (Table 2).

**Figure 2:**
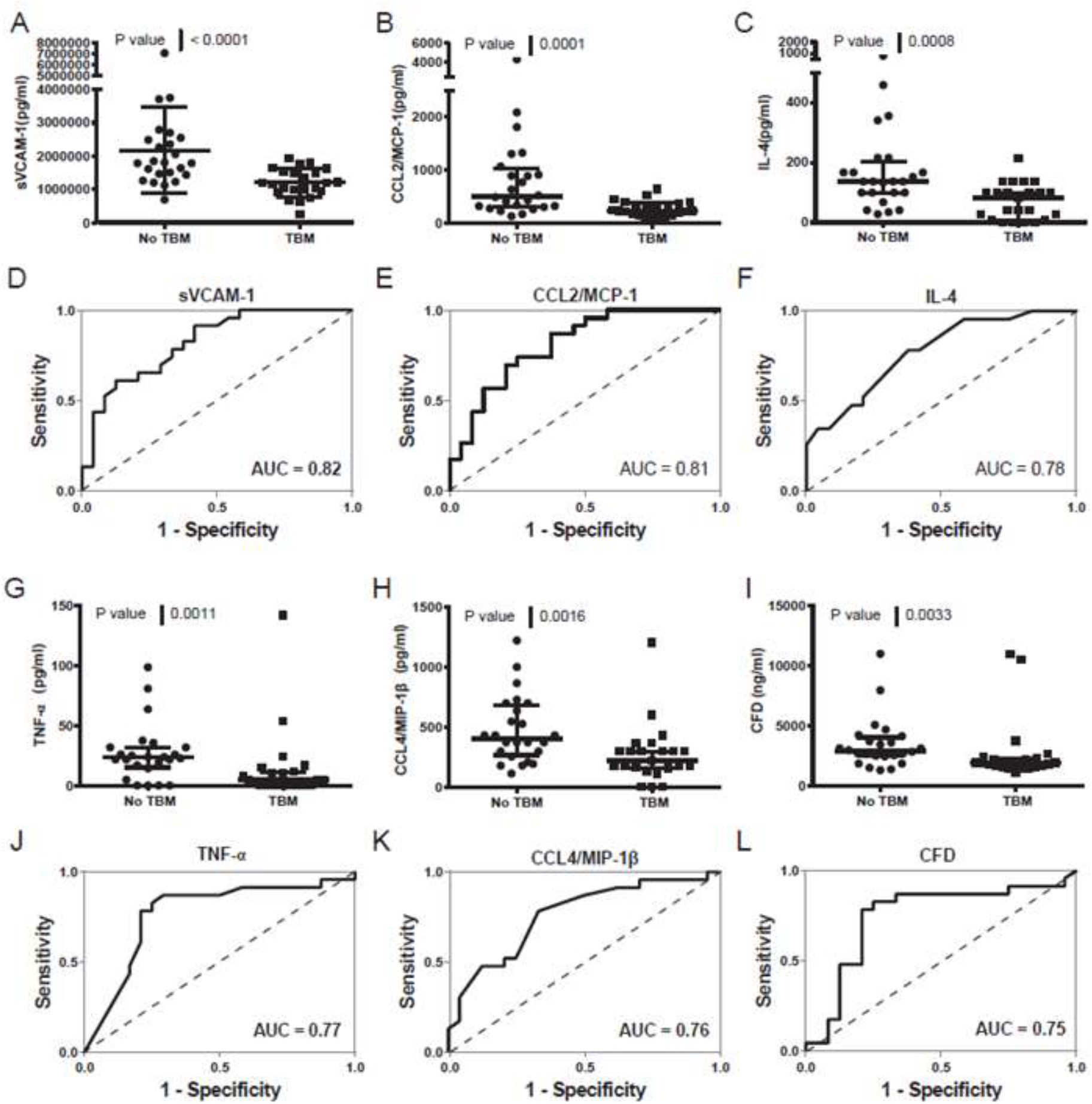
Representative plots showing the concentrations of individual biomarkers detected in serum samples from children with and without TBM and ROC curves showing the accuracies of these biomarkers in the diagnosis of TBM. Error bars in the scatter-dot plots indicate the median and inter-quartile ranges. Representative plots for six analytes with AUC ≥0.75 are shown. The accuracies of all host biomarkers evaluated in the study are shown in Supplementary Table 1.

**Table 2:**
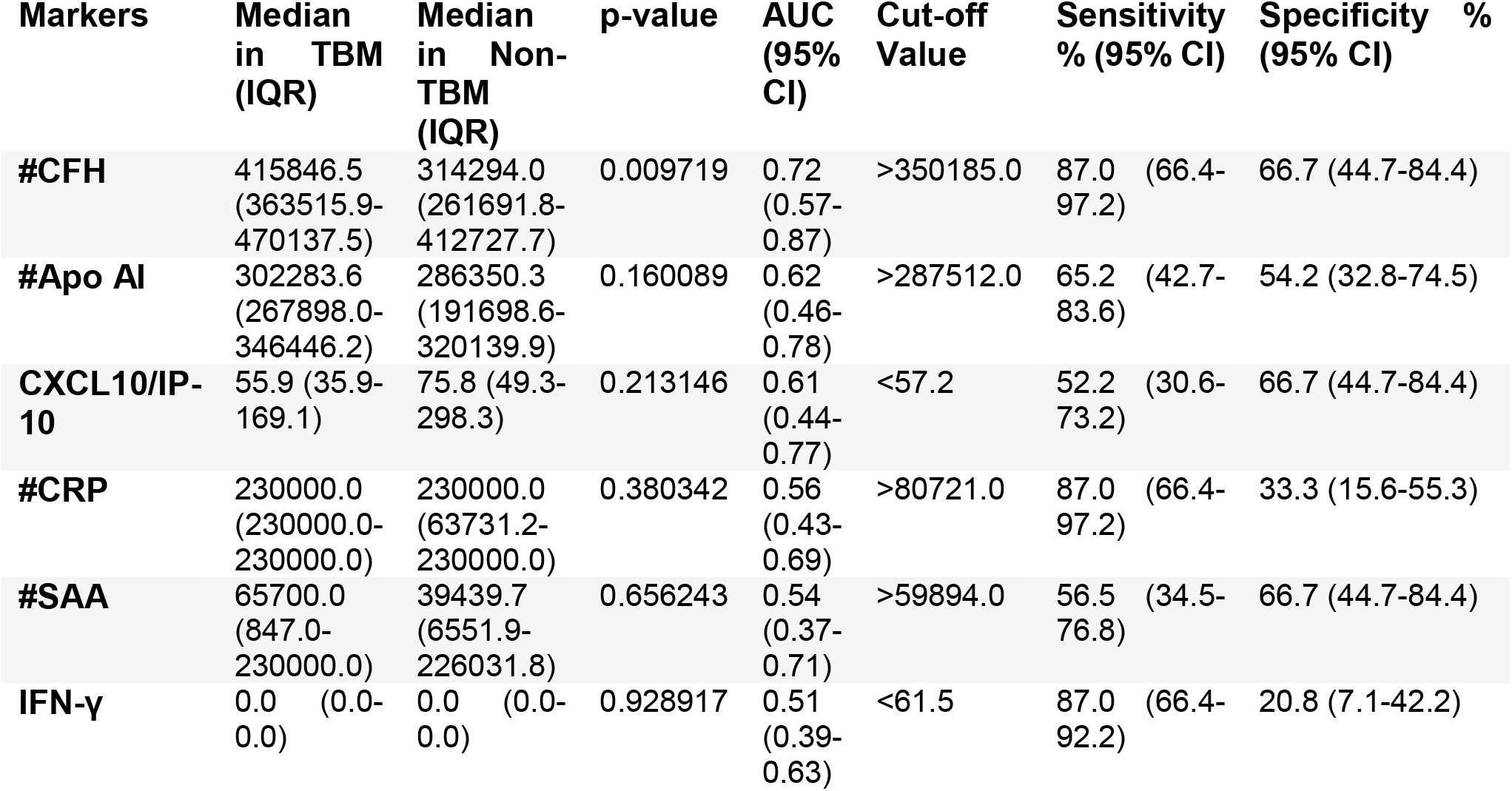
Usefulness of analytes comprising the previously established adult 7-marker serum protein biosignature in the diagnosis of TBM in children. Median levels of host markers detected in serum samples from children with TBM or no TBM (Inter-quartile range in parenthesis) and accuracies in the diagnosis of TBM. Cut-off values and associated sensitivities and specificities were selected based on the Youden’s index. #values shown are in ng/ml, values for all other host markers are in pg/ml.

When these six biomarkers were assessed in combination, the AUC for diagnosis of TBM was 0.75 (95% CI, 0.61-0.90); corresponding to sensitivity of 69.6% (95% CI, 47.1-86.8%) and specificity of 62.5% (95% CI, 40.6-81.2%)). After leave-one-out cross validation, the six-marker combination diagnosed TBM with sensitivity of 65.2% (95% CI, 42.7-83.6%) and specificity of 54.2% (95% CI, 32.8-74.5%). Given that transthyretin, a key biomarker in the seven-marker biosignature was not available for evaluation in this study, NCAM-1 was used as replacement, based on a significant positive correlation with transthyretin (r2 =0.72, p<0.0001) in previous data (27). The modified seven-marker serum protein biosignature including NCAM-1 as replacement for transthyretin predicted TBM with AUC of 0.80 (95% CI, 0.67-0.92); corresponding to sensitivity of 73.9%, (95% CI, 51.6-89.8%) and specificity of 66.7%, (95% CI, 44.7-84.4%). After leave-one-out cross validation, the 7-marker biosignature diagnosed TBM with sensitivity of 60.9% (95% CI, 38.5-80.3%) and specificity of 58.3% (95% CI, 36.6-77.9%), with positive and negative predictive values of 58.3% (95% CI, 44.1-71.4) and 60.9% (95% CI, 45.8-74.1), respectively.

### Utility of other single host serum protein biomarkers in the diagnosis of TBM

When the concentrations of the 63 other host biomarkers (including NCAM-1) were compared between children with and those without TBM using the Mann Whitney U test, the median levels of 16 including sVCAM-1, CCL2, IL-4, TNF-α, CCL4, adipsin, SAP, CC5, G-CSF, IL-10, Apo-CIII, IL-17A, PAI-1, PDGF AB/BB, MBL and NCAM-1 were significantly different (p<0.05) between the two groups, with differences in the concentrations of five markers (CC4b, MMP-1, CXCL8, CC4, sRAGE) showing trends for differences between the two groups (0.05<p≤0.09). The concentrations of SAP, CC5, Apo-CIII, PAI-1, PDGF-AB/BB and MBL were significantly higher in samples from children with TBM whereas those of sVCAM-1, CCL2, IL-4, TNF-α, CCL4, adipsin, G-CSF, IL-10, IL-17A and NCAM-1 were higher in samples from children without TBM (Supplementary table 1). When the diagnostic potentials of individual serum biomarkers were assessed by ROC curve analysis, 13 of these markers showed promise as ascertained by AUC ≥ 0.70 (Supplementary table 1, Figure 3). When only HIV uninfected children were considered, there were improvements in the performances of other host markers including MMP-1 and IL-7 whereas median levels of six including IL-10, MBL, sRAGE CC4, CC4b and NCAM-1 were no longer significantly different (Data not shown).

**Figure 3:**
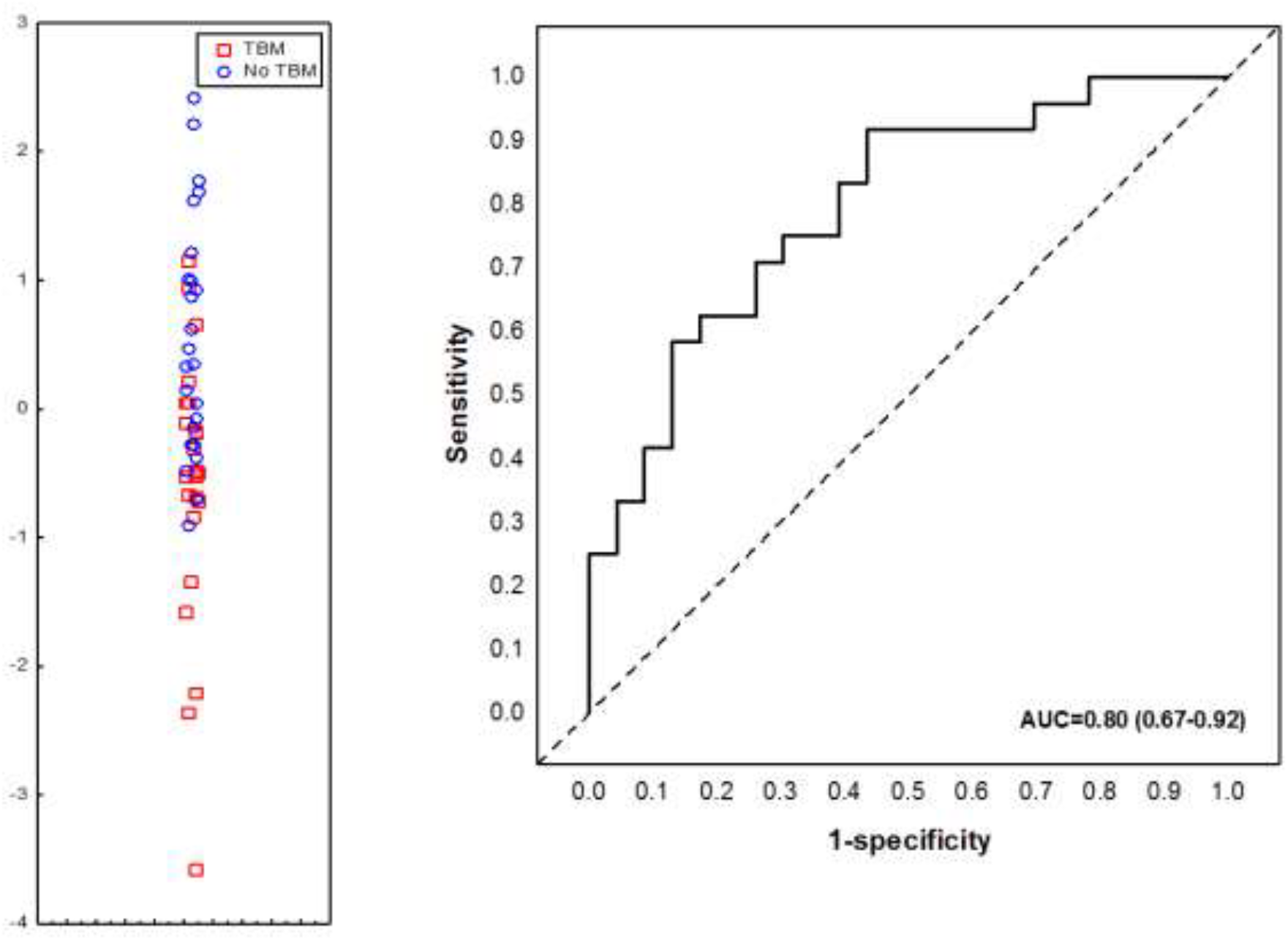
Accuracy of the modified 7-marker serum protein biosignature (CRP, IFN-γ, IP-10, CFH, Apo-A1, SAA and NCAM1) in the diagnosis of TBM in children. Scatter plot showing the ability of the 7-marker signature to classify children as TBM or no TBM (Left image). ROC curve showing the accuracy of the 7-marker biosignature (Right image). Transthyretin in the original 7-marker signature (1) was replaced with NCAM1 as the levels of the two proteins correlated well in a previous study (2) (r^2^ = 0.72; p<0.0001). Red squares; children with TBM; blue circles: children with No TBM.

### Utility of new combinations of host biomarkers in the diagnosis of TBM

When the data obtained from all study participants were fitted into the General Discriminant Analysis (GDA) models regardless of HIV status and regardless of whether biomarkers were part of the adult 7-marker signature or not, optimal prediction of TBM was shown to be achieved with a combination of three markers. The most accurate three-marker biosignature comprising of adipsin, Aβ42 and IL-10 diagnosed TBM with AUC of 0.84 (95% CI, 0.73-0.96); corresponding to a sensitivity of 82.6% (95% CI, 61.2-95.0%) and specificity of 75.0% (95% CI, 53.3-90.2%) (Figure 4). After leave-one-out cross validation, there was no change in the sensitivity of the 3-marker biosignature, whereas the specificity fell to 70.8% (95 CI, 48.9-87.4%). The positive and negative predictive values of the biosignature were 73.1% (95% CI, 58.6-83.9) and 81.0% (95% CI, 62.7-91.5), respectively after leave-one-out cross validation.

**Figure 4:**
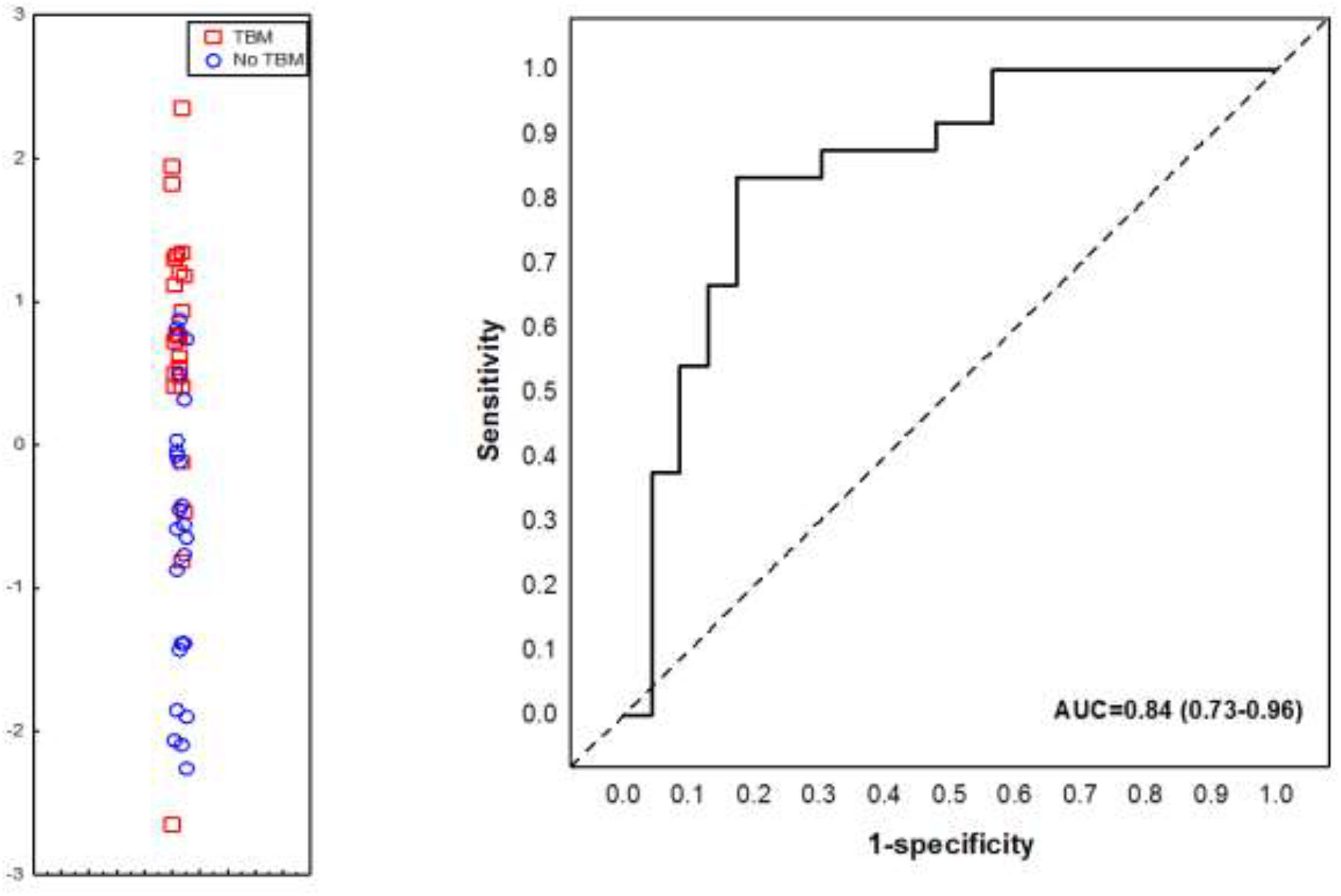
Accuracy of the 3-marker serum biosignature (Complement factor D/adipsin, Ab42 and IL-10) in the diagnosis of childhood TBM. Scatter plot showing the ability of the 3-marker signature to classify children as TBM or no TBM (Left image). ROC curve showing the accuracy of the 3-marker biosignature (Right image). Red squares: children with TBM; blue circles: children with No TBM.

## Discussion

We assessed the usefulness of a modified version of a previously identified adult pulmonary TB 7-marker serum protein biosignature (CRP, IFN-γ, IP-10, CFH, Apo-A1, SAA and NCAM-1, in place of transthyretin) and other host biomarkers that have shown potential in the diagnosis of TB disease in recent adult studies, as tools for diagnosis of TBM in children. Although the modified adult 7-marker serum protein biosignature showed potential as a diagnostic tool for TBM in children, we identified a novel smaller TBM-specific 3-marker serum signature (adipsin, Aβ42 and IL-10), which diagnosed childhood TBM with promising accuracy. It is well-known that most of the poor outcomes resulting from being diagnosed with TB are related to difficulties in the diagnosis of the disease and/or delayed initiation of treatment, especially in high burden settings. The currently available diagnostic tests have several shortcomings, especially in young children and in people presenting with extrapulmonary TB, including TBM (10, 11). The diagnosis of TBM is particularly challenging in resource-constrained areas owing to the expensive and invasive modalities required in making a proper diagnosis. These tests perform poorly individually (29,30), with a combination of several modalities including clinical presentation, CSF findings, neuroimaging, evidence of extraneural TB, and where possible, mycobacteriological confirmation, required for diagnosis (34). Access to these tests requires admission to a tertiary hospital in relatively well-resourced environments. Consequently, children who do not have access these facilities, especially those in resource-constrained, mainly rural settings, miss the opportunity for early diagnosis. Children living in relatively better-resourced areas such as in the Western Cape Province of South Africa still present at the clinic an average of six times prior to eventual proper diagnosis of TBM (20). These factors emphasize the urgent need for new diagnostic tools in childhood TBM.

Host inflammatory protein biomarkers have been shown to possess potential in the diagnosis of TB disease in both adults and children in previous studies (23, 25, 26), including being translated into field-friendly point-of-care tests (20, 21). In a previous study done in our research group, a 3-marker CSF biosignature (VEGF, IL-13 and LL-37) was identified which showed potential as a tool for the diagnosis of TBM in children (25). In a follow-up study, we validated this biosignature but importantly, showed that replacement of two of the proteins in this signature (IL-13 and LL-37) with new proteins (IFN-γ and MPO) resulted in improved accuracy (29). Despite the potential shown by these CSF-based biosignatures, collection of CSF, an invasive procedure that requires lumbar puncture, is challenging in resource-constrained areas. Given the promise shown by blood-based inflammatory biosignatures in pulmonary TB, it is important that similar approaches be assessed for the diagnosis of TBM in children, given that blood will be easier to collect and the fact that blood-based tests may be easily converted into finger prick-based tests as its currently being done in another project (www.screen-tb.eu). In the current study, we have shown proof of principle that blood-based protein biosignatures may be useful in the diagnosis of childhood TBM. Given the limited performance of the adult seven-marker signature in the present study, tests that are developed on adult pulmonary TB patients (e.g. the ScreenTB study) may not perform optimally in children, especially those with TBM as demonstrated in this study. We have however, identified a novel childhood TBM specific 3-marker biosignature which may be considered for further development into a blood-based point-of-care or bedside test for childhood TBM after further validation studies.

A test that is based on blood-based biomarkers such as the biosignature identified in the present study needs to be optimized during future validation and development stages as a screening test for TBM in children. Such a blood-based test is a high-priority need, as identified in the published WHO target product profiles for new diagnostics (35). Further development of such a test into a finger prick blood-based assay will enable easier implementation at the point-of-care or bedside, including in resource-poor settings and may lead to significant reductions in costs, unnecessary lumbar puncture procedures, and the delays that are currently incurred in the diagnosis of TBM in children (36) and consequently, a reduction in the morbidity and mortality currently resulting from TBM. The more invasive and expensive modalities currently being used in the diagnosis of the disease could then only be applied in triaged children with positive point-of-care tests. However, this will only be possible after further validation experiments and incorporation of the biomarkers into such point-of-care test platforms. It is well-known that host inflammatory biomarker-based tests may not be specific for TB, owing to the expression of the biomarkers in other inflammatory conditions and cancer (26). However, it is believed that these specificity concerns may be addressed through the combination of different host biomarkers as done in the present study, with the resultant tests being implemented as triage tests. Given the importance and urgency to commence anti-tuberculous treatment in TBM, a diagnostic test requires adequate sensitivity as well as specificity. It is postulated that a point-of-care test for extrapulmonary TB in adults and children, regardless of HIV co-infection, should have minimum sensitivity of 60% and specificity of 95% when compared to case-defined TBM (38). Both the adult seven-marker serum TB biosignature and our novel smaller TBM-specific 3-marker serum signature had reasonable sensitivity and specificity, with the novel smaller TBM-specific serum signature performing better. It is postulated that the lower specificity than sensitivity of the novel smaller TBM-specific 3-marker serum signature allows it to be considered as a rule-in point of care test. In the correct clinical setting, a rule-in test may lead to early anti-tuberculous treatment initiation, and avoidance of an inevitable deleterious outcome in childhood TBM.

One of the proteins that formed part of our new TBM-specific 3-marker serum protein biosignature for TBM diagnosis is a long form of Amyloid-β (known as Aβ42). Elevated levels of Aβ42 in plasma samples correlate with increased risk of a neurodegenerative disease known as Alzheimer’s disease. However, these levels decrease as disease progresses, supporting the concept of higher accumulation of Aβ42 in neuronal deposits (39). In our study, levels of serum Aβ42 did not show statistical difference between children with TBM and those without TBM. However, these protein contributed to the performance of our TBM-specific 3-marker serum protein biosignature. The association of Aβ42 and dementia in neurodegenerative disease, its decreased levels in bacterial meningitis (40), and its contribution to our signature might suggest its involvement in the pathology of TBM. IL-10 (which also formed part of signature) is a well-known anti-inflammatory and regulatory cytokine. The production of IL-10 is highly increased in healthy human neonates, making this population to be susceptible to TBM (41). Similarly, serum levels of IL-10 were higher in children without TBM in our study. Adipsin/ complement factor D play a role as a regulator in the activation of the alternative complement pathway (42).

The main limitation of the current study was the relatively small sample size, especially individuals with confirmed alternative diagnoses. However, as this study included only children with signs and symptoms suggestive of meningitis, the design of the study was relatively strong and the number of participants enrolled into the study is consistent with the patient numbers that were described in multiple previous studies. Future validation studies should include larger numbers of study participants with suspected meningitis, including those who are HIV infected, and more individuals with confirmed alternative meningitis. This would allow proper validation of the promising signatures by using training and test sets of samples. The use of different biosignatures as tools for triaging children enrolled at lower levels of the healthcare system into those with potentially serious conditions such as meningitis, and other less concerning conditions still needs to be investigated and was not the aim of the present study. Biosignatures, as identified in the current study, will make the greatest impact only if incorporated into field-friendly tests such as finger-prick based assays, for example, based on lateral flow technology. A similar multi-biomarker finger-prick blood-based test is currently under development for the diagnosis of adult pulmonary TB disease (www.screen-tb.eu).

In conclusion, we have shown that a modified version of a previously identified adult 7-marker serum protein biosignature (CRP, SAA, complement factor H, IFN-γ, IP-10, Apo AI and NCAM-1, in place of transthyretin) may be useful in the diagnosis of TBM in children. Furthermore, we identified a smaller childhood TBM specific 3-marker biosignature (adipsin, Aβ42 and IL-10) with a strong potential as a diagnostic tool for childhood TBM. Our findings require validation in larger cohort studies before these biosignatures may be considered candidates for incorporation into point-of-care tests for the early diagnosis of TBM in children.

## Author Contributions

NC and GW conceptualised and designed the study and put together the study team, analysed and interpreted data, and critically revised the manuscript. RS recruited all study participants, analysed and interpreted data, and critically revised the manuscript. KS designed and managed the study database, contributed to data analysis, and revised the manuscript. CM processed the samples in the laboratory, contributed to data acquisition, analysis and interpretation and drafted the manuscript. CS, processed the samples in the laboratory, contributed to data acquisition, and revised the manuscript. All authors provided approval for publication of the content, and agreed to be accountable for all aspects of the work.

## Acknowledgments

We are grateful to all our study participants and support staff for their contribution to this study.

## Data availability

The raw data supporting the conclusions of this manuscript will be made available by the authors, without undue reservation, to any qualified researcher.

