## Supplementary Table 1 for "Potential of host serum protein biosignatures in the diagnosis of tuberculous meningitis in children"

**Supplementary table 1: Utility of alternative host biomarkers detectable in serum samples in the diagnosis of TBM in children.** Median levels of host markers detected in serum samples from children with TBM or no TBM disease (Inter-quartile range in parenthesis) and accuracies in the diagnosis TBM**.** The data shown are raw, untransformed values. Cut-off values and associated sensitivities and specificities were selected based on the Youden's index. *values shown are the absorbance and not the concentration values. #values shown are in ng/ml, values for all other host markers are in pg/ml.

| Markers | Median in TBM (IQR) | Median in Non-TBM (IQR) | p-value | AUC (95% CI) | Cut-off Value | Sensitivity % (95% CI) | Specificity % (95% CI) |
|---|---|---|---|---|---|---|---|
| VCAM-1 | 1197300.0 (952940.0-1543500.0) | 1802000.0 (1456400.0-2521350.0) | 0.000188 | 0.82 (0.70-0.94) | <1580000.0 | 78.3 (56.3-92.5) | 66.7 (44.7-84.4) |
| MCP-1/CCL2 | 244.3 (165.5-390.9) | 512.3 (319.7-994.8) | 0.000262 | 0.81 (0.69-0.93) | <327.3 | 73.9 (51.6-89.7) | 75.0 (53.3-90.2) |
| IL-4 | 82.7 (7.42-99.6) | 136.8 (99.6-191.3) | 0.001147 | 0.78 (0.65-0.91) | <116.7 | 78.3 (56.3-92.5) | 62.5 (40.6-81.2) |
| TNF-α | 4.8 (0.0-11.4) | 23.3 (15.7-31.9) | 0.001457 | 0.77 (0.62-0.91) | <12.9 | 78.3 (56.3-92.5) | 79.2 (57.9-92.9) |
| MIP-1β/ CCL4 | 219.0 (158.1-296.8) | 401.4 (275.7-667.2) | 0.002148 | 0.76 (0.62-0.90) | <334.3 | 78.3 (56.3-92.5) | 66.7 (44.7-84.4) |
| #Adipsin (CFD) | 1950.4 (1611.1-2319.1) | 2917.4 (2493.4-3938.5) | 0.004065 | 0.75 (0.59-0.90) | <2393.0 | 78.3 (56.3-92.5) | 79.2 (57.9-92.9) |
| #SAP | 331539.9 (261542.1-655100.2) | 167660.5 (88309.6-286067.7) | 0.005664 | 0.74 (0.59-0.89) | >257478.0 | 78.3 (56.3-92.5) | 70.8 (48.9-87.4) |

| | | | | | | | |
|---|---|---|---|---|---|---|---|
| #CC5 | 52307.0 (44989.9-59967.4) | 38538.0 (28210.6-47089.8) | 0.006660 | 0.73 (0.58-0.88) | >46742.0 | 69.6 (47.1-86.8) | 75.0 (53.3-90.2) |
| #CFH | 415846.5 (363515.9-470137.5) | 314294.0 (261691.8-412727.7) | 0.009719 | 0.72 (0.57-0.87) | >350185.0 | 87.0 (66.4-97.2) | 66.7 (44.7-84.4) |
| G-CSF | 14.0 (0.0-117.6) | 147.6 (25.1-463.4) | 0.010573 | 0.72 (0.57-0.86) | <76.0 | 65.2 (42.7-83.6) | 70.8 (48.9-87.4) |
| IL-10 | 0.0 (0.0-4.1) | 8.1 (0.0-21.2) | 0.011193 | 0.70 (0.56-0.85) | <7.0 | 95.7 (78.1-99.9) | 54.2 (32.8-74.5) |
| #Apo CIII | 151289.3 (130100.8-181642.4) | 95825.1 (63481.7-161543.8) | 0.014822 | 0.71 (0.55-0.87) | >114926.0 | 87.0 (66.4-97.2) | 62.5 (40.6-81.2) |
| IL-17A | 0.0 (0.0-0.0) | 0.0 (0.0-18.4) | 0.018640 | 0.65 (0.53-0.76) | <11.3 | 95.7 (78.1-99.9) | 37.5 (18.8-59.4) |
| PAI-1(total) | 348736.6 (261199.3-456794.4) | 246289.2 (175941.2-350988.5) | 0.018694 | 0.70 (0.55-0.85) | >255621.0 | 78.3 (56.3-92.5) | 58.3 (36.6-77.9) |
| PDGF-AB/BB | 49576.6 (33649.3-83528.9) | 33592.0 (14786.0-49751.6) | 0.032444 | 0.68 (0.53-0.84) | >42307.0 | 65.2 (42.7-83.6) | 66.7 (44.7-84.4) |
| #MBL | 9533.4 (4686.1-30439.6) | 3299.1 (901.1-14882.4) | 0.033928 | 0.68 (0.52-0.84) | >4522.0 | 78.3 (56.3-92.5) | 58.3 (36.6-77.9) |
| NCAM-1 | 246692.5 (164329.5-305706.5) | 285446.4 (256271.6-342048.0) | 0.036064 | 0.68 (0.52-0.84) | <264419.0 | 69.6 (47.1-86.8) | 70.8 (48.9-87.4) |
| #CC4b | 29843.2 (21128.5-42752.7) | 25562.6 (17752.8-31264.4) | 0.056822 | 0.66 (0.51-0.82) | >26285.0 | 69.6 (47.1-86.8) | 54.2 (32.8-74.5) |

| | | | | | | | |
|---|---|---|---|---|---|---|---|
| MMP-1 | 5694.6 (3233.2-7609.0) | 4084.8 (2174.5-6345.7) | 0.068827 | 0.66 (0.50-0.81) | >4282.0 | 60.9 (38.5-80.3) | 54.2 (32.8-74.5) |
| CXCL8/IL-8 | 37.1 (15.5-54.1) | 55.4 (27.5-112.9) | 0.072101 | 0.65 (0.49-0.81) | <42.1 | 60.9 (38.5-80.3) | 66.7 (44.7-84.4) |
| #CC4 | 157528.9 (90929.3-209684.1) | 85388.5 (48405.5-194821.2) | 0.079129 | 0.65 (0.49-0.81) | >89484.0 | 78.3 (56.3-92.5) | 54.2 (32.8-74.5) |
| sRAGE | 855.2 (773.7-896.6) | 875.8 (855.2-937.8) | 0.094181 | 0.64 (0.48-0.80) | <875.8 | 73.9 (51.6-89.8) | 50.0 (29.1-70.9) |
| TGF-α | 60.3 (26.9-96.2) | 28.5 (5.6-79.8) | 0.110002 | 0.64 (0.48-0.80) | >29.9 | 69.6 (47.1-86.8) | 54.2 (32.8-74.5) |
| IL-7 | 36.0 (22.9-55.8) | 29.2 (12.4-37.7) | 0.110363 | 0.64 (0.48-0.80) | >27.5 | 69.6 (47.1-86.8) | 50.0 (29.1-70.9) |
| IL-6 | 6.8 (1.6-14.6) | 8.9 (2.5-44.7) | 0.135692 | 0.63 (0.47-0.79) | <8.0 | 56.5 (34.5-76.8) | 58.3 (36.6-77.9) |
| GM-CSF | 0.0 (0.0-0.0) | 0.0 (0.0-0.0) | 0.158099 | 0.57 (0.40-0.73) | <9.3 | 100.0 (85.2-100) | 16.7 (4.7-37.4) |
| #Apo AI | 302283.6 (267898.0-346446.2) | 286350.3 (191698.6-320139.9) | 0.160089 | 0.62 (0.46-0.78) | >287512.0 | 65.2 (42.7-83.6) | 54.2 (32.8-74.5) |
| VEGF | 152.4 (112.5-251.2) | 106.7 (74.8-235.8) | 0.169862 | 0.62 (0.45-0.78) | >111.2 | 78.3 (56.3-92.5) | 54.2 (32.8-74.5) |
| Aβ40 | 0.0 (0.0-0.0) | 0.0 (0.0-0.0) | 0.171006 | 0.54 (0.37-0.71) | <72.1 | 100.0 (85.2-100.0) | 8.3 (1.0-27.0) |

| | | | | | | | |
|---|---|---|---|---|---|---|---|
| #CF1 | 66236.8 (49972.0-99204.5) | 54181.1 (45646.3-71882.7)) | 0.176578 | 0.62 (0.45-0.78) | >57835.0 | 65.2 (42.7-83.6) | 62.5 (40.6-81.2) |
| MMP-7 | 808.0 (524.4-1584.1) | 1175.0 (625.5-3399.8) | 0.189921 | 0.61 (0.45-0.78) | <869.0 | 60.9 (38.5-80.3) | 62.5 (40.6-81.2) |
| #Myoglobin | 9.6 (4.4-20.3) | 21.4 (4.9-51.0) | 0.201135 | 0.61 (0.44-0.78) | <10.2 | 60.9 (38.5-80.3) | 66.7 (44.7-84.4) |
| CXCL10/IP-10 | 55.9 (35.9-169.1) | 75.8 (49.3-298.3) | 0.213146 | 0.61 (0.44-0.77) | <57.2 | 52.2 (30.6-73.2) | 66.7 (44.7-84.4) |
| PDGF-AA | 8538.7 (5683.1-15788.5) | 6995.0 (2635.5-12806.3) | 0.221073 | 0.61 (0.44-0.77) | >6150.0 | 69.6 (47.1-86.8) | 50.0 (29.1-70.9) |
| #MIP4 | 241.6 (172.5-366.9) | 178.1 (119.2-342.3) | 0.221073 | 0.61 (0.44-0.77) | >187.7 | 69.6 (47.1-86.8) | 54.2(32.8-74.5) |
| Aβ42 | 0.0 (0.0-0.0) | 0.0 (0.0-556.9) | 0.240593 | 0.58 (0.45-0.72) | <278.4 | 73.9 (51.6-89.8) | 41.7 (22.1-63.4) |
| #CC3 | 40885.9 (36448.0-74127.5) | 46059.4 (25390.4-53871.9) | 0.254876 | 0.40 (0.23-0.57) | >32056.0 | 91.3 (72.0-98.9) | 41.7 (22.1-63.4) |
| #A1AT | 18729.1 (14631.0-24621.2) | 16819.0 (11711.3-27780.9) | 0.287284 | 0.59 (0.42-0.76) | >17908.0 | 60.9 (38.5-80.3) | 58.3 (36.6-77.9) |
| #P-Selectin | 194.3 (102.1-352.1) | 119.1 (54.4-274.0) | 0.330420 | 0.58 (0.42-0.75) | >159.1 | 65.2 (42.7-83.6) | 62.5 (40.6-81.2) |
| CC5a | 2663.1 (1751.2-3946.9) | 2423.2 (1559.1-3554.3) | 0.349063 | 0.58 (0.41-0.75) | >2660.0 | 52.2 (30.6-73.2) | 66.7 (44.7-84.4) |

| | | | | | | | |
|---|---|---|---|---|---|---|---|
| IL-12/23p40 | 0.0 (0.0-0.0) | 0.0 (0.0-0.0) | 0.349077 | 0.52 (0.35-0.69) | <620.1 | 100.0 (85.2-100.0) | 4.2 (0.1-21.1) |
| IL-1β | 0.0 (0.0-0.0) | 0.0 (0.0-9.2) | 0.358788 | 0.56 (0.43-0.68) | <8.3 | 91.3 (72.0-98.9) | 29.2 (12.6-51.1) |
| MMP-8 | 24763.5 (12747.3-86623.8) | 19342.7 (9257.2-35601.6) | 0.360001 | 0.59 (0.41-0.75) | >22769.0 | 56.5 (34.5-76.8) | 58.3 (36.6-77.9) |
| #CRP | 230000.0 (230000.0-230000.0) | 230000.0 (63731.2-230000.0) | 0.380342 | 0.56 (0.43-0.69) | >80721.0 | 87.0 (66.4-97.2) | 33.3 (15.6-55.3) |
| CCL3/MIP-1β | 48.7 (0.0-65.1) | 49.8 (0.0-209.1) | 0.382647 | 0.57 (0.41-0.74) | <48.9 | 65.2 (42.7-83.6) | 54.2 (32.8-74.5) |
| MMP-9 | 205449.19 (59802.48-556493.88) | 174486.7 (73396.4-266465.9) | 0.387093 | 0.57 (0.40-0.74) | >189764.0 | 56.5 (34.5-76.8) | 58.3 (36.6-77.9) |
| IL-21 | 0.0 (0.0-0.0) | 0.0 (0.0-15.9) | 0.396732 | 0.55 (0.43-0.67) | <34.6 | 95.7 (78.1-99.9) | 20.8 (7.1-42.2) |
| Cathepsin D | 439708.5 (308272.3-728466.0) | 493856.4 (331662.9-959098.3) | 0.412583 | 0.57 (0.40-0.74) | <459422.0 | 60.9 (38.5-80.3) | 54.2 (32.8-74.5) |
| ICAM-1 | 216547.6 (137559.5-286618.4) | 215566.5 (171273.1-337326.2) | 0.418679 | 0.57 (0.40-0.72) | <224039.0 | 56.5 (34.5-76.8) | 50.0 (29.1-70.9) |
| #CC9 | 3295.9 (2497.1-4084.6) | 3657.9 (2600.8-4489.9) | 0.475866 | 0.56 (0.39-0.73) | <3502.0 | 65.2 (42.7-83.6) | 58.3 (36.6-77.9) |
| MPO | 4746700.0 (1779300.0-6026200.0) | 3438600.0 (1669250.0-4934150.0) | 0.475891 | 0.56 (0.39-0.73) | >4650000.0 | 52.2 (30.6-73.2) | 70.8 (48.9-87.4) |

| | | | | | | | |
|---|---|---|---|---|---|---|---|
| CD40L | 11633.0 (8228.9-16525.6) | 10742.2 (6930.7-17042.3) | 0.475891 | 0.56 (0.39-0.73) | >11151.0 | 65.2 (42.7-83.6) | 54.2 (32.8-74.5) |
| #GDF-15 | 1.0 (0.6-1.6) | 1.1 (0.6-3.1) | 0.501871 | 0.56 (0.39-0.73) | <1.1 | 60.9 (38.5-80.3) | 54.2 (32.8-74.5) |
| #D-dimer | 9287.6 (1772.3-17900.1) | 9102.9 (3021.2-41007.9) | 0.550286 | 0.55 (0.38-0.72) | <9451.0 | 52.2 (30.6-73.2) | 50.0 (29.1-70.9) |
| BDNF | 15636.7 (10109.5-24406.5) | 18107.0 (8952.6-28946.9) | 0.572783 | 0.55 (0.38-0.72) | <17211.0 | 65.2 (42.7-83.6) | 54.2 (32.8-74.5) |
| CXCL9/MIG | 2309.5 (0.0-3311.4) | 1800.7 (0.0-3557.3) | 0.625319 | 0.54 (0.38-0.71) | >2114.0 | 52.2 (30.6-73.2) | 62.5 (40.6-81.2) |
| #SAA | 65700.0 (847.0-230000.0) | 39439.7 (6551.9-226031.8) | 0.656243 | 0.54 (0.37-0.71) | >59894.0 | 56.5 (34.5-76.8) | 66.7 (44.7-84.4) |
| IL-13 | 0.0 (0.0-338.1) | 0.0 (0.0-756.3) | 0.681743 | 0.53 (0.38-0.69) | <74.6 | 56.5 (34.5-76.8) | 45.8 (25.6-67.2) |
| #CC2 | 15903.9 (8706.0-31171.1) | 15768.9 (6992.3-49343.7) | 0.725481 | 0.53 (0.36-0.70) | <15990.0 | 52.2 (30.6-73.2) | 50.0 (29.1-70.9) |
| Ferritin | 52841.0 (14202.0-114067.7) | 62740.4 (16776.2-169542.0) | 0.740490 | 0.53 (0.36-0.70) | <56314.0 | 56.5 (34.5-76.8) | 58.3 (36.6-77.9) |
| #PEDF | 21756.5 (18654.6-25542.3) | 21401.6 (18159.9-26348.1) | 0.765743 | 0.53 (0.36-0.70) | >21725.0 | 52.2 (30.6-73.2) | 54.2 (32.8-74.5) |
| RANTES | 108231.8 (53485.5-169473.6) | 92692.2 (39285.6-188178.1) | 0.790226 | 0.52 (0.35-0.69) | >99016.0 | 56.5 (34.5-76.8) | 54.2 (32.8-74.5) |

| | | | | | | | |
|---|---|---|---|---|---|---|---|
| #ADMTS13 | 901.2 (545.3-1092.7) | 874.4 (600.0-1120.0) | 0.823096 | 0.52 (0.35-0.68) | <962.3 | 60.9 (38.5-80.3) | 45.8 (25.6-67.2) |
| #NGAL | 394.1 (152.5-1046.1) | 380.5 (189.3-560.3) | 0.831299 | 0.52 (0.35-0.69) | >371.5 | 52.2 (30.6-73.2) | 50.0 (29.1-70.9) |
| CCL1/I-309 | 15.0 (8.6-33.4) | 15.2 (7.6-44.4) | 0.848035 | 0.52 (0.35-0.69) | <15.2 | 52.2 (30.6-73.2) | 50.0 (29.1-70.9) |
| GDNF | 136.3 (120.1-152.7) | 136.3 (136.3-152.7) | 0.921886 | 0.51 (0.34-0.67) | <140.4 | 52.2 (30.6-73.2) | 41.7 (22.1-63.4) |
| IFN-γ | 0.0 (0.0-0.0) | 0.0 (0.0-0.0) | 0.928917 | 0.51 (0.39-0.63) | <61.5 | 87.0 (66.4-92.2) | 20.8 (7.1-42.2) |
| *Cathelicidin-LL37 | 0.5 (0.3-0.9) | 0.5 (0.3-0.9) | 0.974533 | 0.49 (0.31-0.66) | >0.4 | 60.9 (38.5-80.3) | 34.8 (16.4-57.3) |
| S100B | 2800.0 (2744.2-2800.0) | 2800.0 (2744.2-2800.0) | 0.986591 | 0.50 (0.34-0.66) | >2772.0 | 55.6 (30.8-78.5) | 40.0 (19.1-64.0) |
